# Human-caused ignitions and plant invasion drive fire-regime change in a dryland biodiversity hotspot

**DOI:** 10.64898/2026.11.03.764038

**Authors:** Kyle C. Rodman, Adam L. Mahood, Anita J. Antoninka, Matthew A. Bowker, Helen I. Rowe

## Abstract

Dryland ecosystems cover much of Earth’s terrestrial surface and are increasingly affected by climate warming, land-use change, and biological invasion. The Sonoran Desert of western North America, a biodiversity hotspot that historically experienced infrequent fire, exemplifies how these global-change processes are driving ecosystem transformation through the emergence of novel disturbance regimes. Using records of more than 73,000 wildland fire events, we quantified anthropogenic and environmental controls on fire ignition, size, and spread across the Sonoran Desert. Human-caused ignitions accounted for 95.5% of all recorded fires since the early 1990s and were concentrated near roads, buildings, and other areas of human activity, particularly during late spring. In contrast, lightning-caused ignitions were rare within the Sonoran Desert and occurred primarily in adjacent montane forests during late summer months. Consistent with broader trends across the western United States, fires were typically largest when they ignited far from roads, where access is limited, and fire management often relies on indirect attack strategies. Pre-fire annual and perennial herbaceous vegetation both promoted fire spread, with detectable effects at cover values as low as 5%, while bare ground reduced fuel connectivity and constrained fire growth. Rather than exhibiting a distinct threshold response, fire spread increased linearly with herbaceous cover, suggesting that even modest increases in fine fuels can alter fire behavior in this historically fuel-limited ecosystem. Together, our results illustrate that human ignitions determine where fires occur, whereas herbaceous fuels determine whether those ignitions spread, driving fire-regime transformation across an iconic North American desert. Managing and adapting to this novel fire regime will require coordinated efforts to reduce human-caused ignitions, limit invasive herbaceous fuels, maintain bare-ground interspaces, and rapidly suppress fires. More broadly, our findings identify the mechanisms controlling fire regime change in dryland ecosystems and provide a framework for anticipating ecosystem transformation under ongoing global change.

## 1. Introduction

Dryland ecosystems cover nearly half of Earth’s terrestrial surface and are threatened by interacting global change drivers, including climate change, shifting human land use, and biological invasions (Lewin et al., 2024; Prăvălie, 2016; Ren et al., 2022; Wang et al., 2022). In combination, these factors are altering processes such as fire regimes, a key determinant of species distributions and community structure, globally (Bowman et al., 2020; Jones et al., 2022; Pausas et al., 2025). It follows that when fire regimes deviate from the conditions under which local populations and communities evolved, rapid ecosystem transformation can occur (Johnstone et al., 2016; Turner & Seidl, 2023). Historically, fire was infrequent in many drylands because fire ignition and spread were limited by sparse and discontinuous fuels (Abatzoglou & Kolden, 2011; Krawchuk & Moritz, 2011; Pausas et al., 2025). However, invasion by non-native herbaceous plants can increase fuel continuity and establish positive feedbacks that promote recurrent burning (Brooks et al., 2004; D’Antonio & Vitousek, 1992). These dynamics have been documented in drylands worldwide, including Australian savannas (Bowman et al., 2014), South African shrublands (Rahlao et al., 2009), and deserts of western North America (Balch et al., 2013; Bradley et al., 2018). Climate variability further enhances dryland flammability because unusually wet years can increase herbaceous fuel production (Crimmins et al., 2025; Littell et al., 2009; Ohlert et al., 2026) while human activities facilitate both plant invasions and fire ignitions (Aplet et al., 2026; Gelbard & Belnap, 2003). Together, these drivers can shift dryland ecosystems from infrequent-fire systems limited by both fuels and ignitions toward increasingly fire-prone states (Fusco et al., 2022).

Despite growing recognition that these interacting processes are increasing fire activity in drylands, important uncertainties remain regarding the mechanisms driving fire-regime change. Human-caused ignitions are increasingly common in many dryland ecosystems, but they are not uniformly distributed across the landscape (Balch et al., 2017; Fusco et al., 2016). Human-caused fires are often concentrated near roads and other infrastructure (Aplet et al., 2026; Syphard et al., 2025), patterns that can be evaluated using spatial records of fire ignitions and their causes (e.g., Short, 2022). Accessibility and proximity to infrastructure can also influence fire management strategies and whether an ignition develops into a large fire event (Baylis & Boomhower, 2023; Thompson et al., 2021; Yocom et al., 2019), relationships that can be evaluated using ignition locations and recorded fire size information (Aplet et al., 2026). Herbaceous fuels enhance fire spread (Archibald et al., 2012; Mahood et al., 2023; Tortorelli et al., 2023), yet the cover required to support active fire behavior and the relative contributions of different species and functional groups remain uncertain in many dryland ecosystems (Brooks et al., 2004; Costanzo & Munson, 2026). Fortunately, annual maps of functional group cover are now available across many regions (e.g., Allred et al., 2021), enabling detailed evaluations of relationships between vegetation cover and fire behavior (Rodman et al., 2023). Although climate is not generally considered a limitation to burning in drylands (Krawchuk & Moritz, 2011), novel fire-weather datasets (e.g., Vitolo et al., 2020) permit evaluation of whether aridity and fuel moisture influence fire ignition and spread. Together, these emerging methods and geospatial datasets provide new opportunities to evaluate how fuels, climate, and human activities interact to shape novel fire regimes. Understanding these interactions is necessary to predict dryland ecosystem vulnerability and identify opportunities for effective management under ongoing global change (Aslan et al., 2021; Chambers et al., 2019; Fusco et al., 2022).

The Sonoran Desert of western North America provides an ideal system for evaluating the mechanisms of fire-regime change in drylands (Fig.1). The region is a globally significant biodiversity hotspot that supports a wide range of vertebrates, arthropods, and plant species, including the iconic saguaro cactus (*Carnegiea gigantea*) (Chávez-Hernández et al., 2026; Albuquerque et al., 2025; Dimmitt, 2000; Orr et al., 2021). A shift toward frequent surface fire would represent a major departure from the evolutionary history of many native species. Before the 20th century, fires were likely small and infrequent (Esque & Schwalbe, 2002; Thomas, 1991; Zouhar, 2023). The human population has more than tripled across this region since the 1980s (Fig. 1b; Pesaresi et al., 2024), fueling significant expansion of the wildland-urban interface (Radeloff et al., 2018). These land-use changes are likely influencing patterns of fire ignitions (Fusco et al., 2016; Madakumbura et al., 2026) and seasonality (Cattau et al., 2020). At the same time, invasive annual grasses, perennial grasses, and forbs are increasing fuel connectivity and flammability (Aslan et al., 2021; Gornish & Howery, 2019; McDonald & McPherson, 2013). Although large fires remain a relatively recent phenomenon in the Sonoran Desert (Fig. 1c), they can have substantial ecological consequences because many native species are poorly adapted to frequent burning (Abella, 2009; Cave & Patten, 1984; Rogers & Steele, 1980; Wilder et al., 2021). Consequently, understanding the factors driving fire-regime transformation in the Sonoran Desert is both an urgent conservation priority and an opportunity to better understand ecosystem transformation across drylands worldwide.

**Figure 1:**
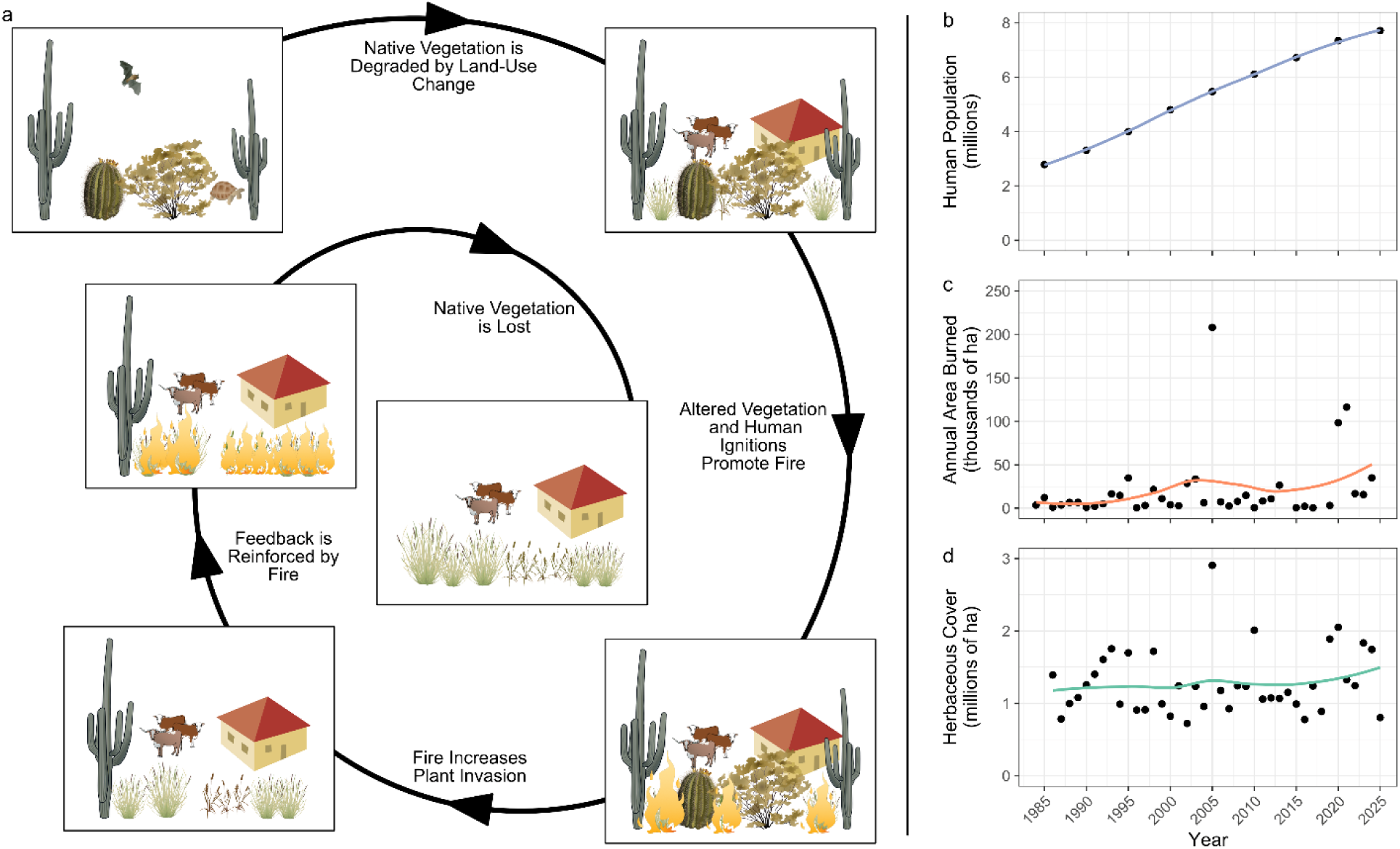
Land-use change and species invasions are altering fire regimes and ecosystems in the Sonoran Desert. Panel (a), adapted from Fusco et al. (2022), shows a typical sequence in which intact Sonoran ecosystems are degraded due to these interacting drivers. Time series of (b) human population, (c) annual area burned in large fires, and (d) herbaceous cover are from the Global Human Settlement Layer (Pesaresi et al., 2024), Monitoring Trends in Burn Severity (Eidenshink et al., 2007), and Rangeland Analysis Platform (Allred et al., 2021) datasets, respectively, within the Sonoran Desert Ecoregion (CEC, 2025). Colored lines in (b-d) show interdecadal trends using loess smooths; all three variables have evidence of positive trends since the 1980s based on Theil-Sen slope estimates. Graphics in (a) are from the University of Maryland Integration and Application Network (https://ian.umces.edu/media-library).

Here, we evaluated how global change drivers shape three key components of fire-regime transformation: fire ignition, fire size, and fire spread. Using geospatial analyses of more than 70,000 fire events across the Sonoran Desert, we addressed three questions: (1) Which factors are most strongly associated with human- and lightning-caused fire ignitions and where are they most likely to occur? (2) Which factors influence whether an ignition becomes a large fire? (3) How does pre-fire cover of different functional groups influence fire spread? By integrating large fire-occurrence datasets with spatial analyses of fire ignition, size, and spread, we quantify the relative importance of fuels, climate, and human activities in shaping contemporary fire regimes in the Sonoran Desert. Further, a spatially explicit investigation of these processes helps to identify opportunities for precise and targeted management under ongoing global change.

## 2. Methods

### 2.1. Study Area and Invasive Species of Concern

We defined our study area using North American Level III Ecoregions from the Commission for Environmental Cooperation (CEC, 2025), including the Sonoran Desert (10.2.2), Madrean Archipelago (12.1.1), and Arizona/New Mexico Mountains (13.1.1). Sonoran Desert vegetation primarily occurs within Ecoregion 10.2.2 but also extends into adjacent uplands to the north and east (Rollins, 2009). Because key data describing fire occurrence (Short, 2022) and functional group cover (Allred et al., 2021) were not consistently available across international boundaries, we restricted analyses to the US portion of the Sonoran. In this area, mean annual temperature is 21.5°C and mean annual precipitation is 196 mm year^-1^, with approximately one-third falling during the North American monsoon (July-September) and much of the remainder occurring in winter and spring (1991-2020 climate normals; *PRISM Climate Group, Oregon State University*, 2026). Invasive plant species of greatest concern include red brome (*Bromus rubens*), Mediterranean grass (*Schismus* spp.), buffelgrass (*Pennisetum ciliare*), fountain grass (*Pennisetum setaceum*), Lehmann’s lovegrass (*Eragrostis lehmanniana*), and stinknet (*Oncosiphon piluliferum*) (Gornish & Howery, 2019; Van Devender & Dimmitt, 2000).

### 2.2. Fire Data, Q1-Q3

For Questions 1 and 2, related to fire ignition and fire size, we used wildfire occurrence records from version 6 of the FPA-FOD database (Short, 2022). These data include fire ignition locations, dates, assigned causes, and final area burned for all known fire records in the United States from 1992 to 2020. We retained occurrences within the Sonoran Desert, Madrean Archipelago, and Arizona/New Mexico Mountains ecoregions, excluding records within US Census-designated urban areas and those with an unknown cause. The resulting dataset included 73,207 fire occurrences. We also used these records to characterize seasonal patterns of human-and lightning-caused ignitions and the causes of human-started fires. We extracted the ignition (or discovery) date, cause, location, and final fire size for use in subsequent analyses.

For Question 3, related to fire spread, we combined fire perimeter data from the Monitoring Trends in Burn Severity (MTBS) program (Eidenshink et al., 2007) and the National Interagency Fire Center (NIFC, 2026). We included all fires with >10 ha core area (after applying a 60-m inward buffer) between 1987 and 2024, removing duplicate records. Earlier fires were excluded because pre-fire land-cover data began in 1986 (Allred et al., 2021). We retained any fires that intersected the Sonoran Desert ecoregion and were fully contained within the broader study area (Sonoran Desert, Arizona/New Mexico Mountains, and Madrean Archipelago), reducing the influence of fires that primarily burned in adjacent forested uplands. Following these restrictions, the final dataset included 346 fires with perimeter information.

### 2.3. Environmental Variables Used to Predict Fire Activity, Q1-Q3

For use in each analysis, we compiled spatial predictors describing vegetation, topography, fire weather, and human infrastructure (Table 1). Vegetation cover variables were obtained from the Rangeland Analysis Platform (Allred et al., 2021), topographic variables from USGS 3DEP elevation models (USGS, 2026), and fire weather metrics from ERA5 reanalysis data (Vitolo et al., 2020). Human-use variables included distances to roads, buildings, transmission lines, mines, and water bodies. Additional descriptions of data acquisition and processing are provided in Appendix S1. We used Google Earth Engine (Gorelick et al., 2017) as well as the packages *terra* (Hijmans, 2025) and *sf* (Pebesma, 2025) in R software (R Core Team, 2024) for all spatial data processing.

**Table 1:**
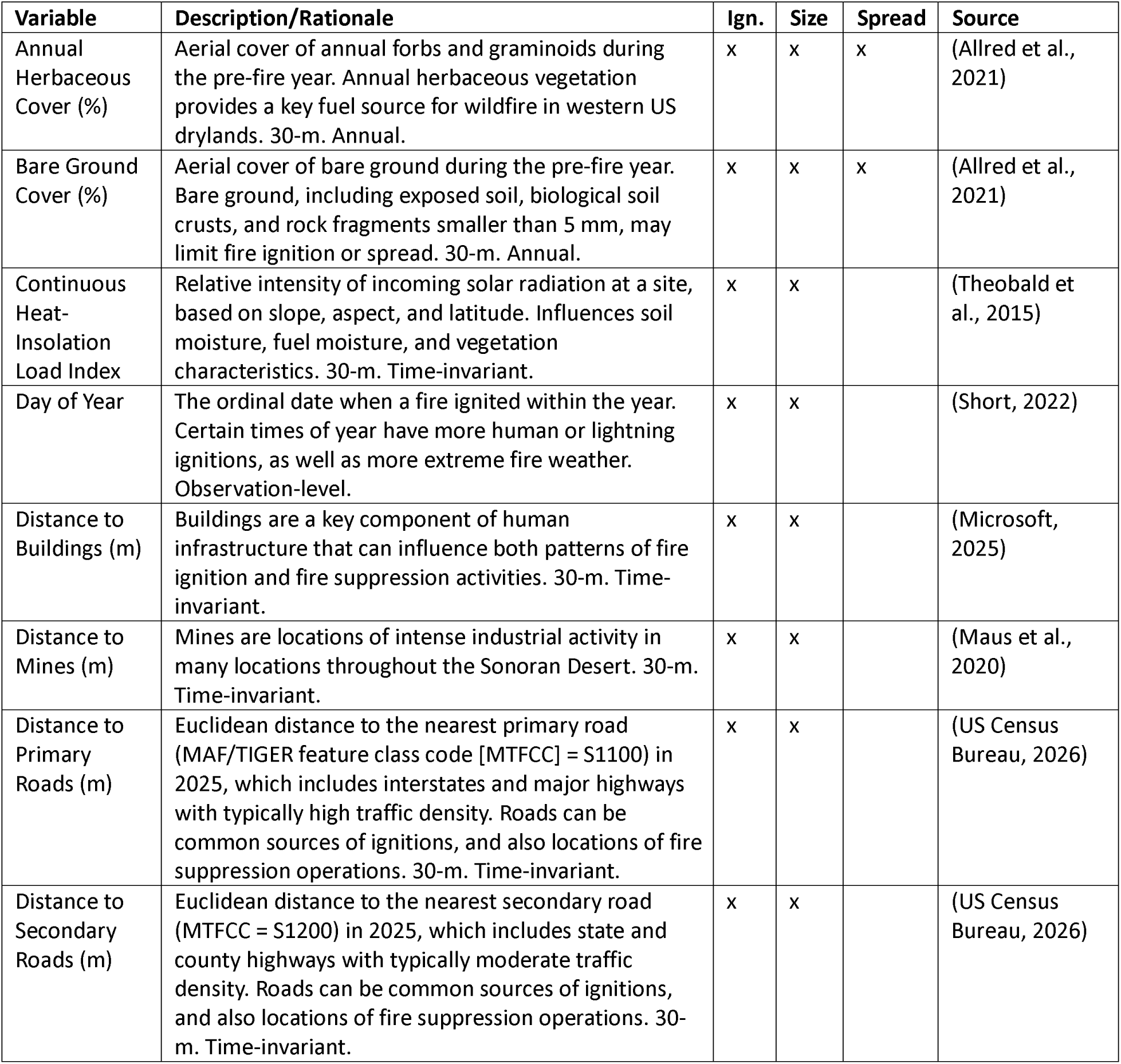
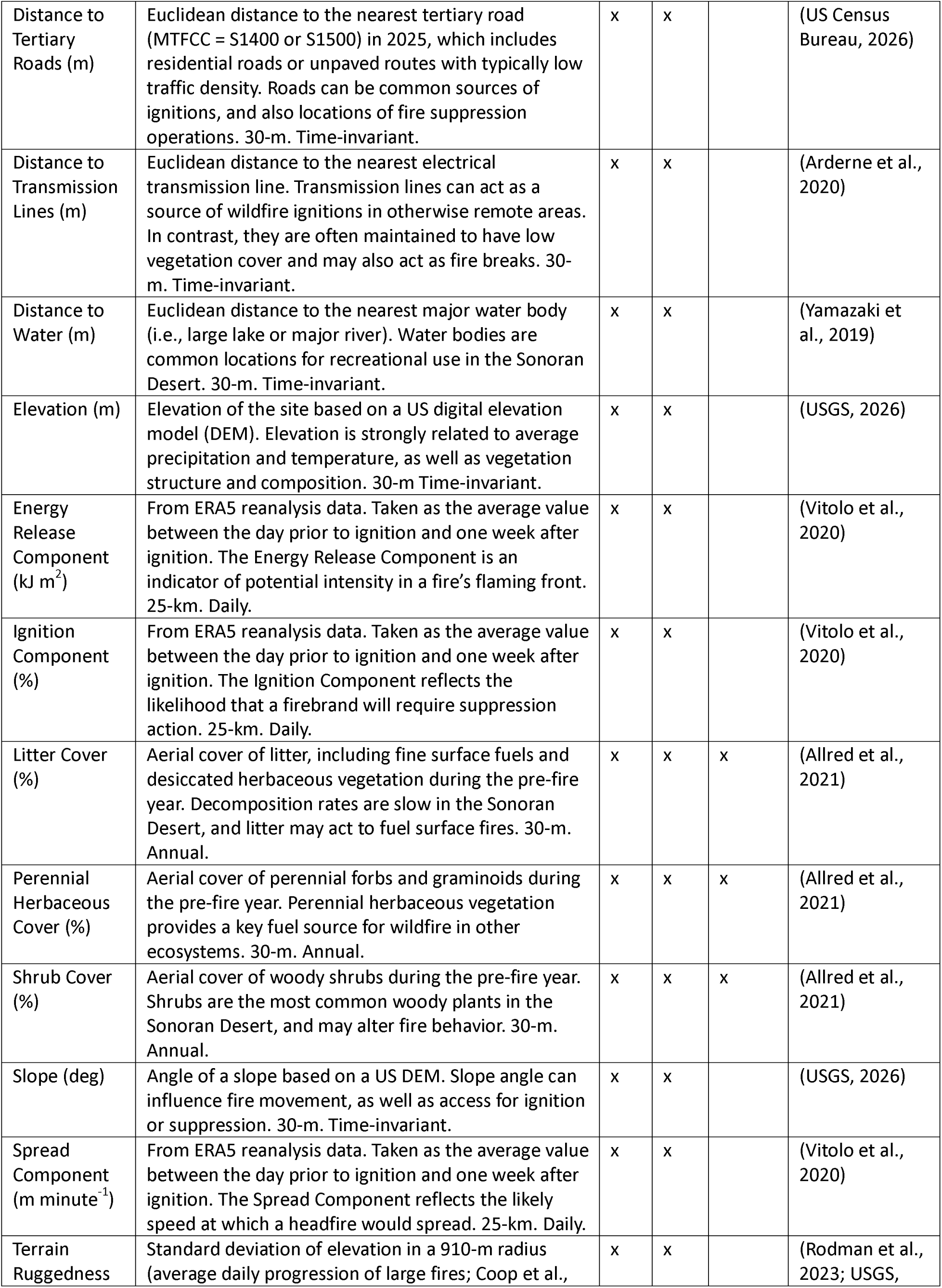
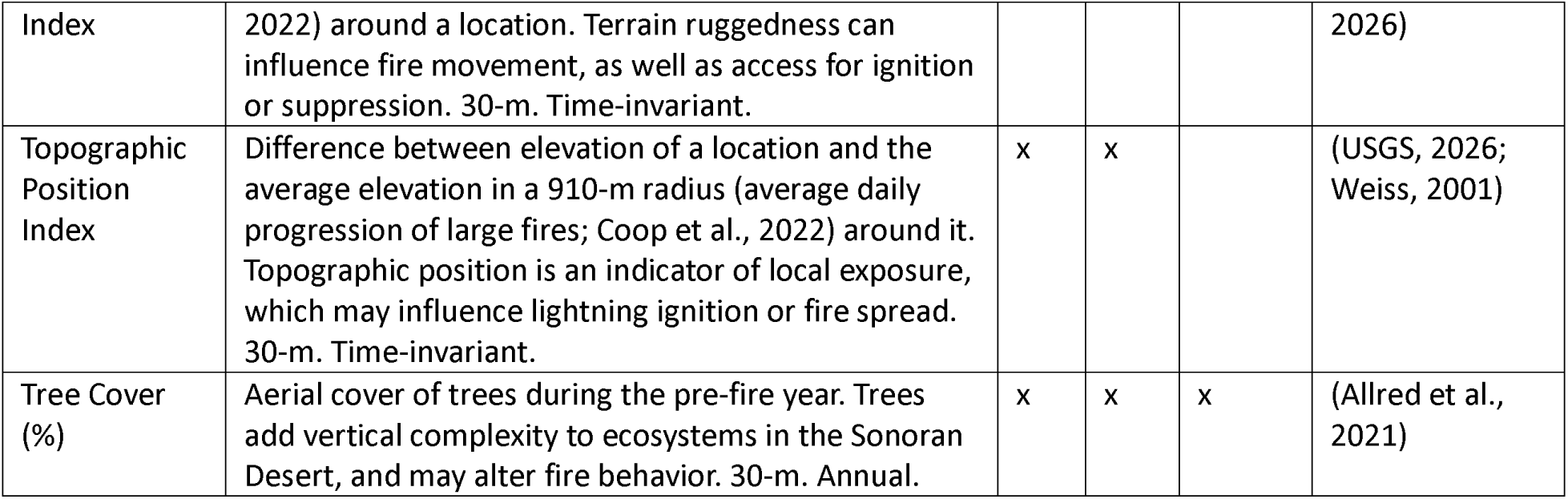
Summary of variables used in analyses of fire ignition (Ign.), size, and spread in the Sonoran Desert. Fire spread used a different modeling framework with only pre-fire cover of six functional groups, whereas ignition and size used all variables as potential covariates.

### 2.4. Statistical Analyses, Q1 and Q2 – Random Forests

We fit three Random Forest (RF) models (Breiman, 2001) to evaluate drivers of (1) human-caused ignitions, (2) lightning-caused ignitions, and (3) fire size. Because ignition records represent presence-only data, we generated 50,000 pseudo-absence points, which were sufficient to characterize background environmental variation across the study area. Ignition models were developed for binary classification (fire occurrence vs. pseudo-absence), whereas fire size was modeled using RF regression of a continuous response. We used a natural-log transformation of fire size to reduce the influence of extreme values. For ignition models, we used a Hellinger split rule, a maximum tree depth of two, and 2,000 trees (Valavi et al., 2021). For fire-size models, we used a maxstat split rule, no limit to tree depth, and 1,000 trees (Wright & Ziegler, 2017).

For each model, we followed the general RF model selection and validation procedures of Rodman et al. (2023). First, we excluded highly collinear predictors using a variance inflation factor threshold of 10; only the Energy Release Component was removed at this stage. We then used 20-fold spatial cross-validation, with observations assigned to spatial blocks using k-means clustering, and forward feature selection to identify predictors for inclusion in final models (Meyer et al., 2018, 2023). A similar cross-validation framework was then used to tune minimum node size and the number of variables evaluated at each split. To reduce overfitting, we increased minimum node size or removed predictors when necessary based on model diagnostics. Next, we evaluated model performance using out-of-bag predictions from final fitted models. To do so, we quantified ignition-model performance using area under the receiver operating characteristic curve (AUC) and fire-size-model performance using Pearson’s correlation between observed and predicted natural-log-transformed fire size. To evaluate sensitivity to locational uncertainty in ignition records (Short, 2022), we repeated all analyses using predictors aggregated to the 1.6-km resolution of the Public Land Survey System grid (Appendix S2).

Using the fitted RF models, we predicted patterns of human-caused ignitions, lightning-caused ignitions, and fire size throughout the study area. We restricted predictions to areas outside of development, agricultural lands, or surface water using the 2021 National Land Cover Dataset (Wickham et al., 2026). To develop final predictive maps, we used 90th-percentile fire-weather conditions, mean ignition dates, 2025 RAP vegetation cover, and the most recent versions of all other predictors (Table 1). Thus, mapped predictions represent the relative likelihood of human and lightning ignitions (Models 1 and 2) and expected fire size (Model 3) under relatively severe fire-weather conditions during the 2026 fire season. We back-transformed fire size predictions by exponentiation such that mapped values represent geometric mean fire size in ha. We used the *ranger* package (Wright & Ziegler, 2017) for all RF model development and the *CAST* (Meyer et al., 2023) and *caret* packages (Kuhn, 2008) for model selection and tuning in R software.

### 2.5. Statistical Analyses, Q3 – Fire Perimeter Effect

The fire perimeter effect (FPE) quantifies the influence of vegetation and landscape characteristics on fire spread and perimeter formation (Harris et al., 2025). FPE is calculated as the normalized difference between pre-fire cover at the fire perimeter (“edge”) and within the fire interior (“core”) using the following equation:

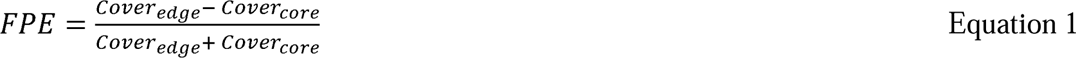

We defined core areas as fire polygons with a 60-m inward buffer and edges as a 120-m band centered on the fire perimeter (60 m inside and outside the mapped perimeter). The resulting index ranges from -1 to 1 and can be compared directly among fires with different levels of pre-fire cover. Negative values indicate that a cover type is associated with increased fire spread or less effective suppression, whereas positive values indicate the opposite. We calculated FPE for all fires that intersected the Sonoran Desert ecoregion and contained at least 10 ha of core area. Thus, we did not include fires that were contained in upland areas of adjacent ecoregions. Using RAP cover data from the year preceding each fire, we calculated FPE separately for annual herbaceous cover, perennial herbaceous cover, litter, shrubs, trees, and bare ground, using mean values of fractional cover in core and edge areas. We also evaluated whether thresholds of annual herbaceous, perennial herbaceous, and total herbaceous cover were associated with changes in the FPE. For each cover type, we created binary maps representing cover above or below thresholds of 5%, 10%, 15%, 20%, 25%, and 30%, calculated proportional cover within fire cores and edges, and converted these values to FPE metrics. For each analysis, we tested whether FPE values differed from zero using two-sided Wilcoxon signed-rank tests with Bonferroni-adjustments for multiple comparisons.

## 3. Results

### 3.1. Sources of Ignitions

Human activities were the dominant source of fire ignitions in the Sonoran Desert Ecoregion between 1992 and 2020, accounting for 95.5% of all reported fires with a known cause. In contrast, lightning-caused ignitions represented just 4.5% of fires in the Sonoran Desert, but the majority (54.0%) of fires in the adjacent Arizona/New Mexico Mountains and Madrean Archipelago Ecoregions were lightning-caused. Human- and lightning-caused ignitions also exhibited distinct seasonal patterns (Fig. 2a). Lightning ignitions peaked during the North American Monsoon in late summer, whereas human-caused ignitions were most common in late spring. Human-caused fires arose from a variety of activities, including escaped campfires, arson, debris burning, and vehicle use (Fig. 2b).

**Figure 2:**
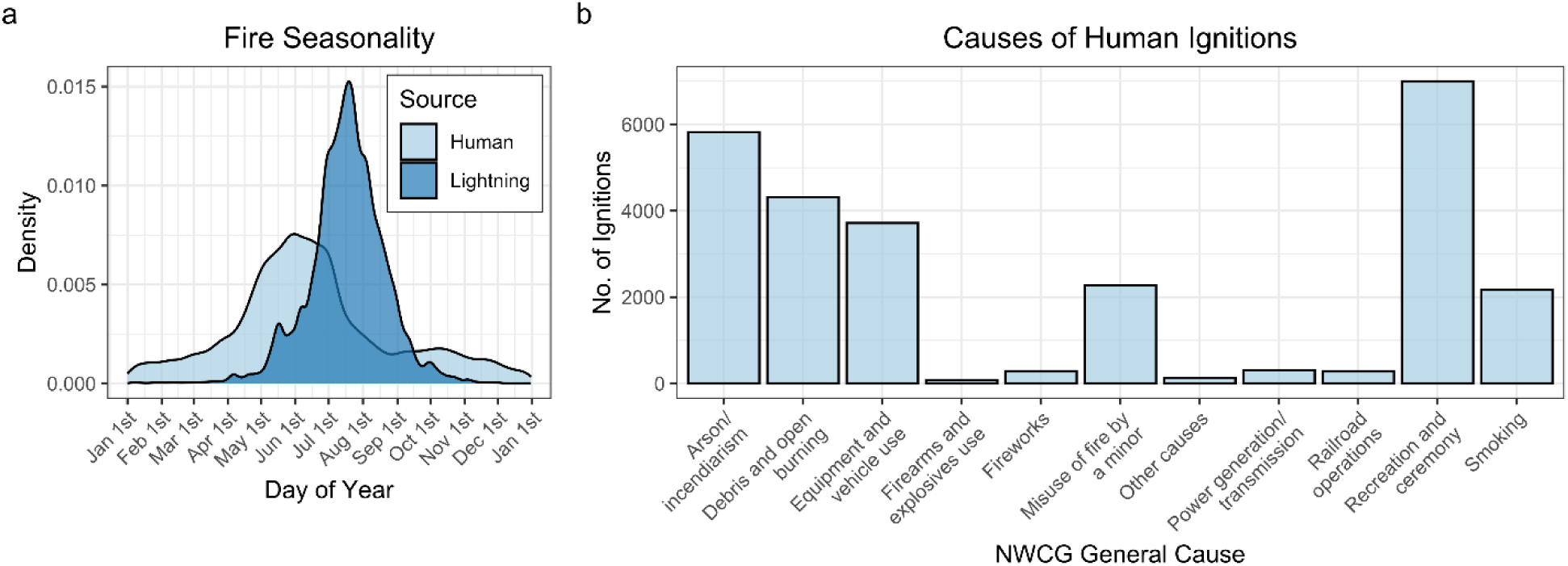
Seasonality of human- and lightning-caused wildfires throughout the study area (a) and reported causes of human ignitions (b). In (a), fire dates were based on the reported date of discovery. In (b), causes were based on the National Wildfire Coordinating Group (NWCG) general-cause category; 17,120 human ignitions with unknown causes were excluded.

In the final Random Forest (RF) model, human-caused ignitions were most strongly associated with patterns of development and infrastructure (Fig. 3). For example, human ignitions were most likely to occur near buildings, transmission lines, and roads (Fig. 3c-f, h). Notably, transmission lines caused just a small fraction of recorded fires (Fig. 2b), suggesting this variable is acting as a proxy for human activity density rather than a direct ignition source. Relative likelihood of ignition also increased when the spread component exceeded approximately 10 m minO¹ (Fig. 3g). Consistent with observed seasonal patterns (Fig. 2a), human ignitions were most common during spring and early summer (i.e., before July 15^th^) (Fig. 3i). Finally, bare ground exhibited a weakly negative relationship with ignition likelihood (Fig. 3j). Regional model predictions reflected these relationships, with the highest relative likelihood of human ignitions occurring near communities and along transportation corridors (Fig. 3b). Model performance was strong, with an out-of-bag AUC of 0.85.

**Figure 3:**
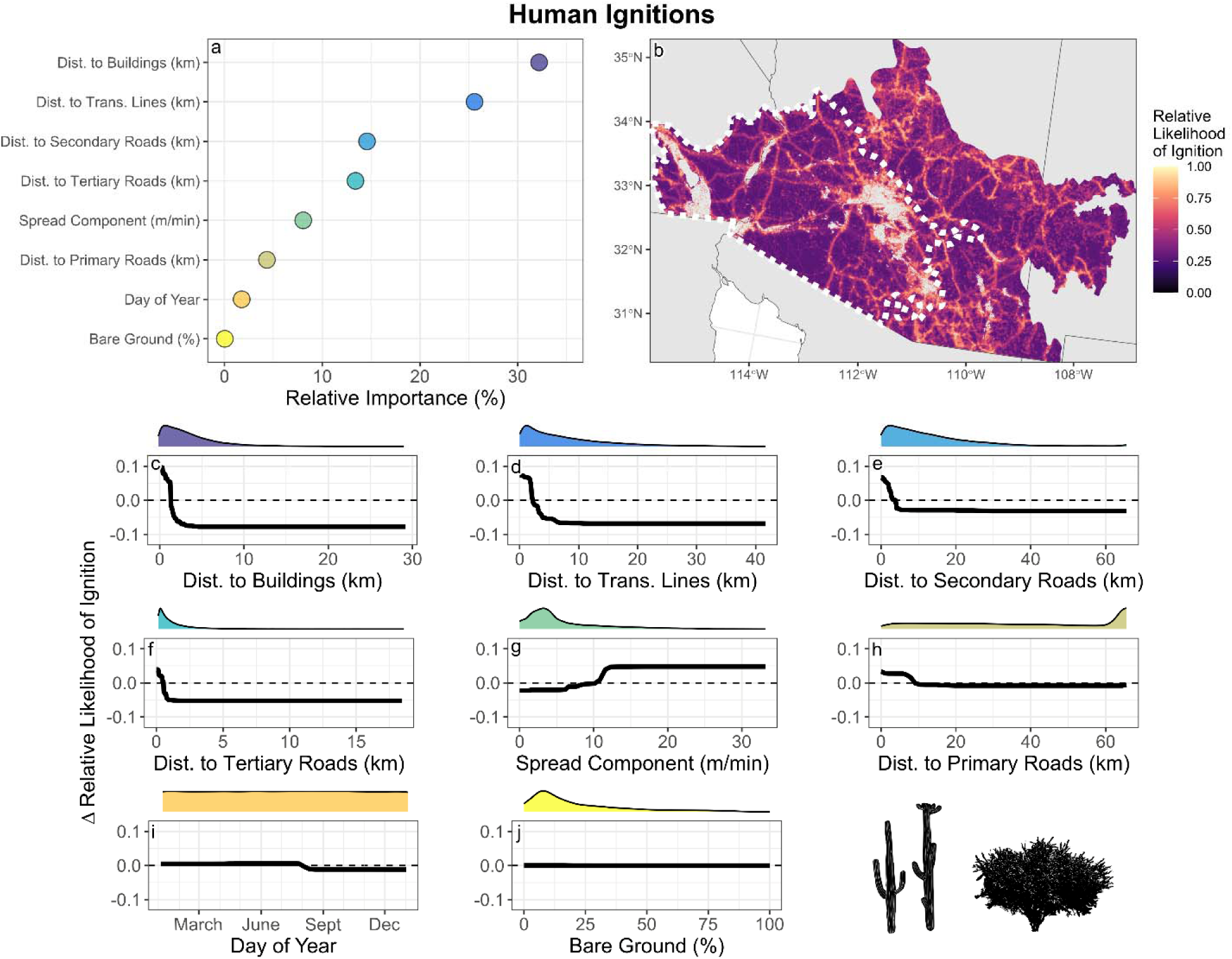
Results of the Random Forest model predicting locations of human-caused ignitions throughout the study area. Variable importance (a) and accumulated local effects (ALE) plots (c-j) show the relative importance of predictors and their effects on model predictions, respectively. Density curves above ALE plots show the distribution of each predictor within the study area. The predictive map (b) was generated using 90^th^-percentile Spread Component values, the mean ignition date for human-caused fires, and observed spatial variation in all other covariates; the dashed white line indicates the extent of the Sonoran Desert Ecoregion within the United States. Icons for saguaro (*Carnegiea gigantea*) and Sonoran palo verde (*Parkinsonia praecox*) were obtained from PhyloPic (https://phylopic.org).

Elevation and seasonality were the strongest predictors of lightning-caused ignitions in the final RF model (Fig. 4). Elevation accounted for more than 50% of total variable importance (Fig. 4a), and lightning-caused ignitions had much greater likelihood above 1,500 m (Fig. 4c). Lightning-caused ignitions were most common during summer months between June and September (Fig. 2a, Fig. 4d). The relative likelihood of lightning-caused ignition declined with increasing bare ground cover (Fig. 4e) and increased with Spread Component (Fig. 4f). Lastly, lightning-caused ignitions exhibited a weakly negative relationship with Ignition Component, suggesting that they were more likely to occur during periods with high fine-fuel moisture (Fig. 4g). Predicted ignition patterns reflected these relationships, indicating that lightning-caused fires were uncommon within the Sonoran Desert Ecoregion but occurred frequently in high-elevation areas of the surrounding ecoregions (Fig. 4b). Model performance was very strong, with an out-of-bag AUC of 0.91.

**Figure 4:**
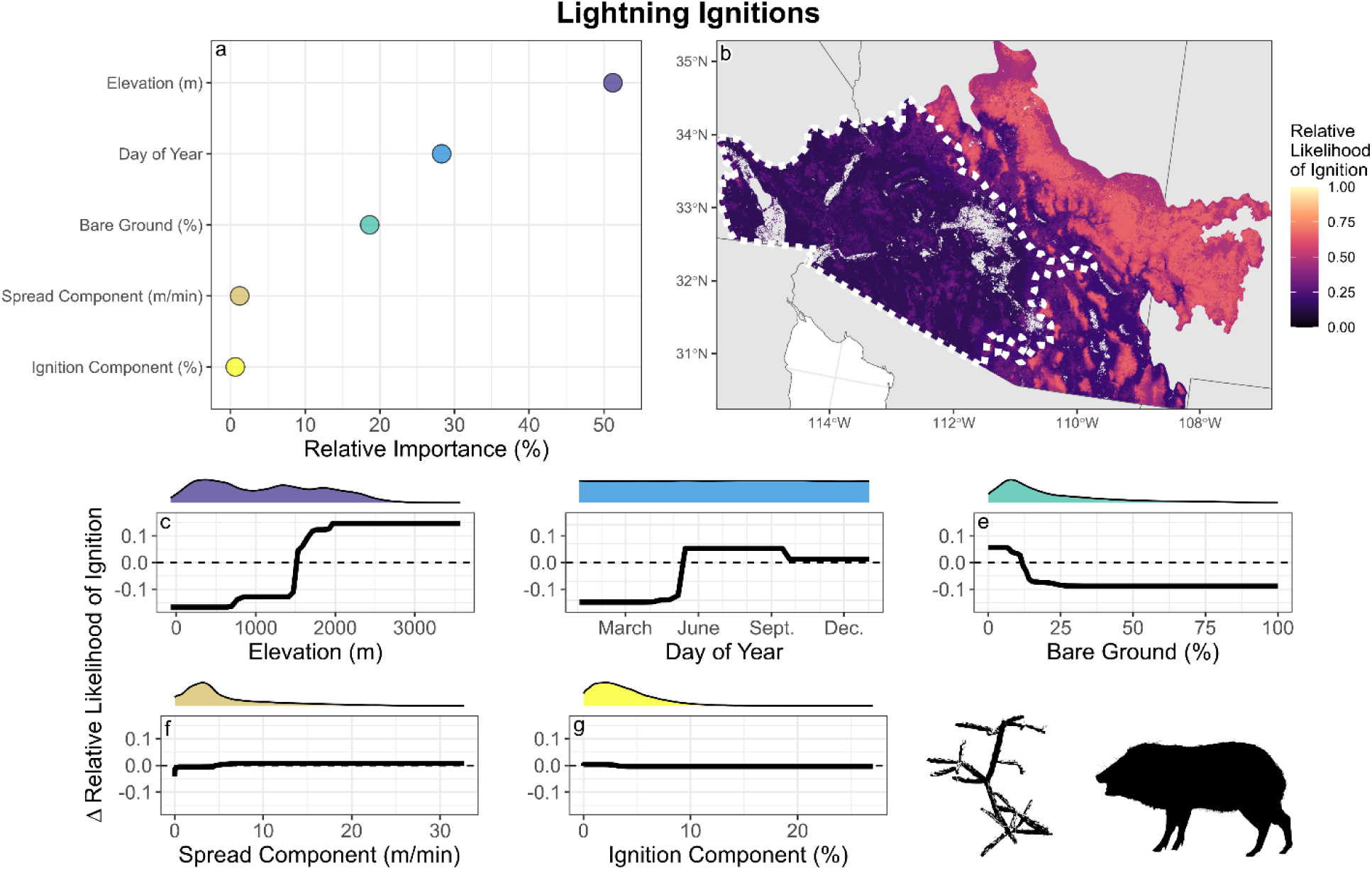
Results of the Random Forest model predicting locations of lightning-caused ignitions throughout the study area. Variable importance (a) and accumulated local effects (ALE) plots (c-g) show the relative importance of predictors and their effects on model predictions, respectively. Density curves above ALE plots show the distribution of each predictor within the study area. The predictive map (b) was generated using 90^th^-percentile values of Ignition and Spread Components, the mean ignition date for lightning-caused fires, and observed spatial variation in all other covariates; the dashed white line indicates the extent of the Sonoran Desert Ecoregion within the United States. Icons for Thurber’s cholla (*Cylindropuntia thurberi*) and javelina (*Dicotyles tajacu*) were obtained from PhyloPic (https://phylopic.org).

### 3.2. Fire Size

Fire size was related to environmental conditions, vegetation, and human infrastructure (Fig. 5). Elevation was the strongest predictor, with fires becoming larger when ignitions occurred below approximately 1,500 m (Fig. 5c). Pre-fire perennial herbaceous cover was also strongly related to fire size, where ignitions in areas with >25% perennial herbaceous cover often led to larger fires (Fig. 5d). Fire size increased under high values of the Ignition Component, indicative of windy, dry conditions and low fine-fuel moisture (Fig. 5e). Larger fires also tended to originate farther from roads (Fig. 5f,h), transmission lines (Fig. 5g), and buildings (Fig. 5i). Predicted fire sizes reflected these relationships, with the largest fires occurring in lower-elevation, remote areas characterized by high perennial herbaceous cover (Fig. 5b). Model performance was modest, with an out-of-bag Pearson’s correlation of 0.34.

**Figure 5:**
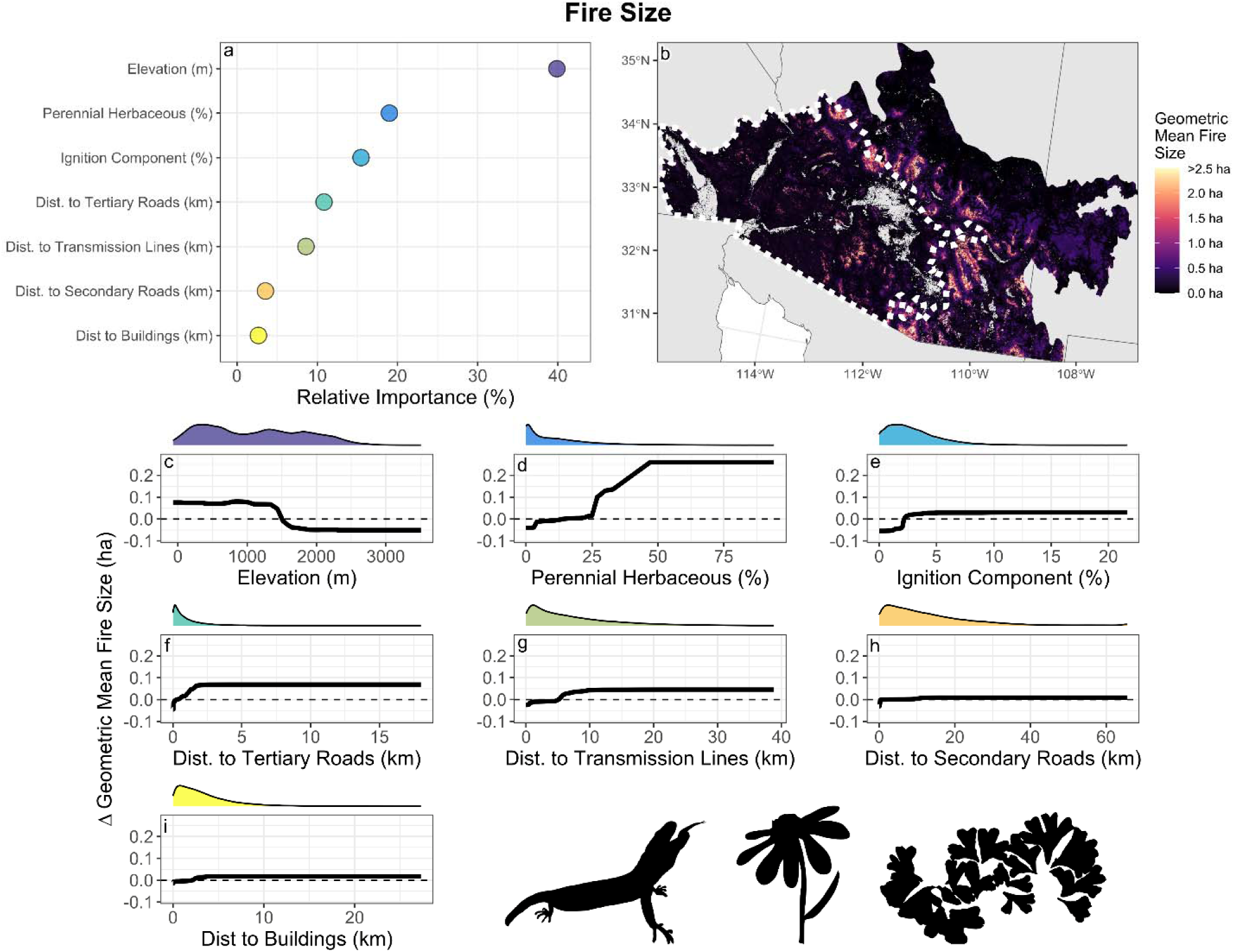
Results of the Random Forest model predicting fire size throughout the study area. Variable importance (a) and accumulated local effects (ALE) plots (c-i) show the relative importance of predictors and their effects on model predictions, respectively. Density curves above ALE plots show the distribution of each predictor within the study area. The predictive map (b) was generated using a 90^th^-percentile value of the Ignition Component and observed spatial variation in all other covariates; the dashed white line indicates the extent of the Sonoran Desert Ecoregion within the United States. Icons for Gila monster (*Heloderma suspectum*), brittlebush (*Encelia farinosa*), and common crystalwort (*Riccia sorocarpa*) were obtained from PhyloPic (https://phylopic.org).

### 3.3. Pre-Fire Cover and Fire Spread

Fire Perimeter Effect (FPE) analyses indicated that fire spread was strongly related to pre-fire vegetation cover across the Sonoran Desert (Fig. 6). Annual and perennial herbaceous cover had the lowest FPE values, indicating that they promoted fire spread relative to other cover types (Fig. 6a). In contrast, bare ground had consistently positive FPE values, indicating that it inhibited fire spread. Litter and tree cover also had significantly negative FPE values, although effect sizes were modest and median values remained near zero. Shrub cover did not differ significantly from zero, suggesting a limited influence on fire spread. Annual and perennial herbaceous vegetation exhibited little evidence of threshold behavior, with median FPE values declining linearly as cover increased (Fig. 6b-d). FPE values at the highest levels of annual or perennial herbaceous cover were not significantly different from zero; however, these cover classes were present in a relatively small number of fires, likely influencing the ability to detect statistical differences. Overall, relationships in both functional groups were consistent with the idea that fire spread increased with greater amounts of herbaceous cover.

**Figure 6:**
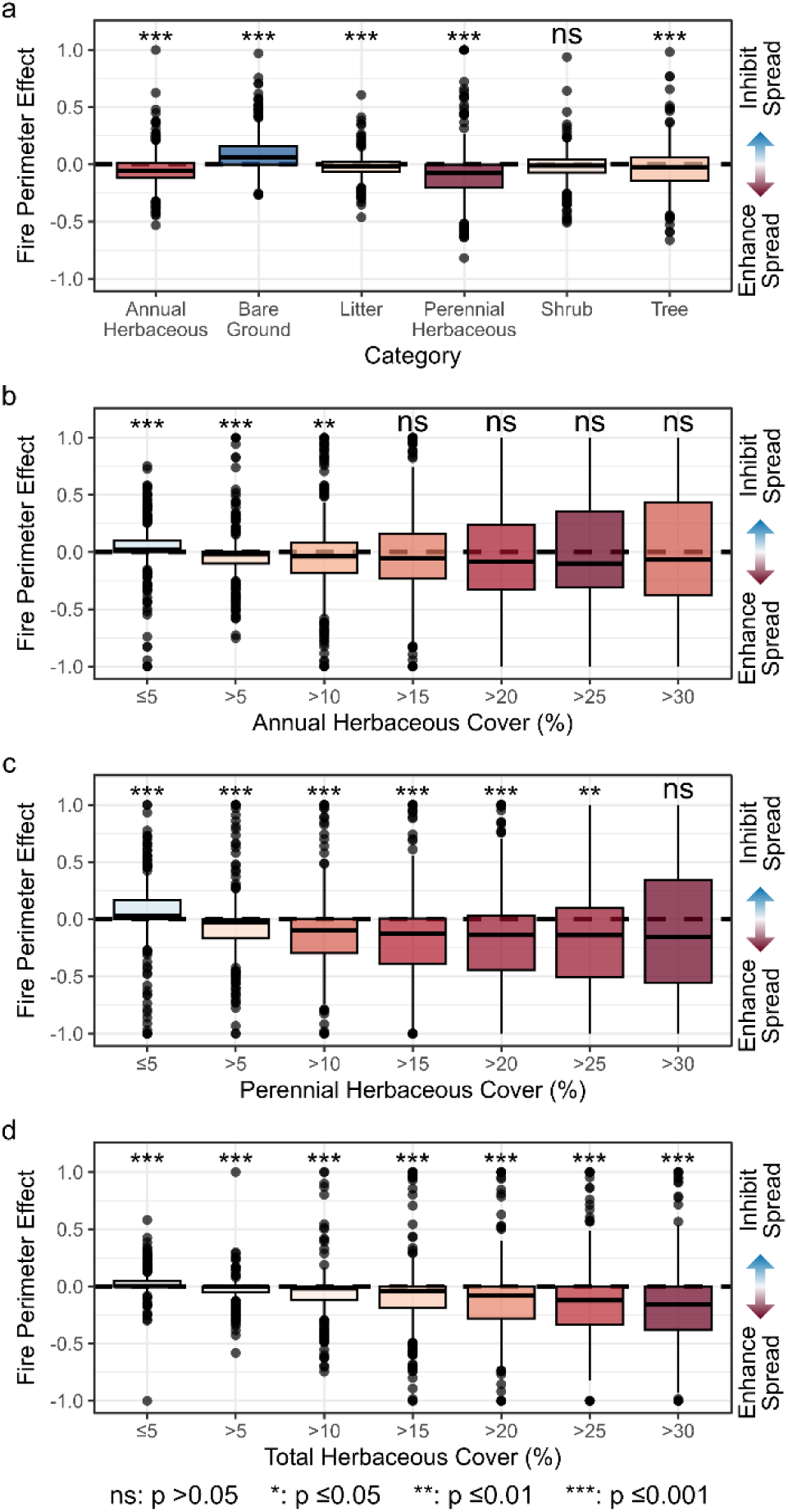
Relationships between pre-fire vegetation cover and fire spread in the Sonoran Desert Ecoregion, based on the Fire Perimeter Effect (Harris et al. 2025). Boxplots show the distribution of fire-level FPE values, where negative values indicate that a cover type facilitates fire spread and positive values indicate that a cover type inhibits fire spread. Box fill colors are based on the absolute value of the median FPE within each panel, with darker colors indicating larger departures from zero. Significance levels indicate whether FPE values differed from zero. In panels (b-d), median FPE values generally declined with increasing herbaceous cover, indicating little evidence of threshold behavior.

## 4. Discussion

Our results demonstrate that contemporary fire activity in the Sonoran Desert is governed by distinct processes that control ignition and spread. Human activities determine where fires occur, whereas herbaceous fuels determine whether they expand and become large. Consistent with this framework, we found that human activities near development and infrastructure were responsible for most fire ignitions in the Sonoran Desert, whereas lightning-caused fires were rare and primarily occurred in upland areas. The largest fires tended to occur in remote areas where fire management access was limited. Lastly, fire size and spread in the Sonoran were strongly promoted by herbaceous cover and limited by bare-ground interspaces. Taken together, these findings indicate that a human-grass-fire cycle has become established in this dryland biodiversity hotspot, and also help to identify opportunities for targeted management, including fire prevention, suppression, and invasive species control. More broadly, our study provides a framework for disentangling how ignitions, fuels, and environmental conditions interact to drive fire-regime transformation in dryland ecosystems worldwide.

### 4.1. Human Activity Drives Fire Occurrence in an Ignition-Limited Ecosystem

Human activities were the source of 95.5% of all fires with a known cause in the Sonoran Desert from 1992-2020 (Short, 2022), and these fires overwhelmingly occurred near roads, buildings, and other infrastructure (Fig. 3). Human-caused fires have broadly increased in the southwestern US over recent decades (Syphard et al., 2025), concurrent with a rapid expansion of the wildland-urban interface (Fig. 1b; Radeloff et al., 2018). While lightning-caused fires are common in upland areas throughout the Southwest (Fig. 4; Balch et al., 2017), they are rare in the Sonoran Desert (Humphrey, 1963). Historical fire activity in these natural-ignition-limited systems probably reflected a combination of infrequent lightning ignitions and localized Indigenous burning practices (Bahre, 1985; Roos et al., 2022; Zouhar, 2023), suggesting that widespread human ignitions represent an unprecedented change to fire regimes across the region. Importantly, this novel ignition pressure is not uniformly distributed across the landscape. We found that buildings, roads, and transmission lines were strongly associated with human ignitions and collectively accounted for more than 90% of total variable importance in the RF model, overwhelming the influence of environmental and climatic predictors. Modern development and infrastructure exert strong controls on fire occurrence across ecosystems worldwide (Aplet et al., 2026; Archibald et al., 2009; Coogan et al., 2020; Dorph et al., 2022). In the Sonoran Desert, these relationships are particularly consequential because human populations and infrastructure now regulate fire occurrence in an ecosystem where natural ignitions were historically rare.

At the same time, ignitions that occurred in remote locations, far from infrastructure, were more likely to become large fires (Fig. 5f-i). Road networks and other infrastructure have complex effects on fire regimes (Aplet et al., 2026; Johnston et al., 2021). In drylands, roads can enhance ignition pressure and facilitate the spread of non-native plants that increase fuel connectivity (Fusco et al., 2022; Gelbard & Belnap, 2003). In contrast, transportation networks can improve access for fire management and are commonly used as containment lines during suppression operations (Thompson et al., 2021; Yocom et al., 2019). Because of these opposing effects, management strategies that limit public access during periods of elevated fire danger (e.g., forest or area closures) are commonly used to reduce ignitions while maintaining access for fire operations (Edgeley et al., 2025). More broadly, debates around road construction and land management, such as the proposed rescission of the 2001 Roadless Area Conservation Rule in the US, highlight trade-offs associated with expanding transportation infrastructure in fire-prone landscapes. Our results suggest that expanded infrastructure may enhance suppression access but also increase ignition pressure and opportunities for non-native plant invasion. Humans are key regulators of both fire ignition and fire growth in dryland systems, shaping how novel fire regimes emerge under ongoing global change.

### 4.2. Herbaceous Vegetation Cover Increases Fire Size and Enhances Spread

Whereas human activities strongly influenced ignition pressure, herbaceous vegetation primarily influenced fire size and spread in the Sonoran Desert (Fig. 5, Fig. 6). Previous research on fire-climate relationships in the western US indicates that antecedent moisture is a major predictor of annual area burned in drylands (Crimmins et al., 2025; Littell et al., 2009) because it promotes the production and continuity of fine fuels (e.g., Ohlert et al., 2026; Shaw et al., 2023). Our results provide additional insight into these relationships, suggesting that vegetation functional groups differ in their influence on fire activity. Perennial herbaceous cover was strongly associated with both fire size and spread (Fig. 5d; Fig. 6), with greater effects as cover increased. Although native perennial grasses occur naturally throughout the Sonoran Desert (Dimmitt, 2000), invasion by non-native perennial species such as buffelgrass and fountain grass has increased fuel continuity, landscape flammability, and potential fireline intensity (Costanzo & Munson, 2026; McDonald & McPherson, 2013; Wilder et al., 2021). Annual species promoted fire spread, but their effects were weaker than those of perennial herbaceous vegetation. This difference may partly reflect the annual temporal resolution of RAP cover data (Allred et al., 2021); winter annuals that emerge and senesce during the same year as a fire would not be fully represented by our pre-fire vegetation estimates. Nevertheless, both perennial and annual herbaceous cover had stronger relationships with fire spread than any other functional group, underscoring the importance of fine-fuel abundance and connectivity to contemporary fire regimes in the Sonoran Desert.

Without management intervention, fire and the expansion of invasive herbaceous species will continue to degrade native-dominated ecosystems in drylands of the western US (Chambers et al., 2019; Mahood & Balch, 2019; Wilder et al., 2021). Although removal of woody vegetation is a stated objective of many fuel-reduction treatments in our study area (Laushman et al., 2020), woody plant cover exhibited only weak relationships with fire size and spread in our analyses (Fig. 5, Fig. 6a). Instead, our results suggest that management of herbaceous fuels may provide the greatest opportunity to reduce fire activity in the Sonoran Desert. We also found that bare-ground interspaces strongly constrained fire spread, indicating that treatments that reduce herbaceous fuel continuity may be particularly effective. Traditional approaches for the treatment of invasive herbaceous species include mechanical removal or herbicide use (pre-emergent or post-emergent), often with multiple treatments (Costanzo & Munson, 2026; Rowe et al., 2023). Restoration of biological soil crusts (biocrusts) represents a complementary approach because intact biocrust communities can stabilize soils, maintain bare-ground interspaces, and limit colonization by invasive plants that promote the human-grass-fire cycle (Bacovcin et al., 2025; Havrilla et al., 2019). Methods to restore biocrusts are under development (Bowker et al., 2025; Rowe et al., 2026), but additional research is needed on how to scale their production and optimize their placement in the field. Protecting fire refugia during suppression operations and accelerating post-fire recovery through targeted revegetation may help limit fire-driven ecosystem transformation and improve the persistence of native plant communities following fire (Hulvey et al., 2017; Meddens et al., 2018; Stevens et al., 2021).

### 4.3. Limitations and Future Research Needs

Several sources of uncertainty should be considered when interpreting our results. Although we analyzed ignition and spatial covariate data at a 30-m resolution, the underlying FPA-FOD occurrence records have a maximum reported horizontal error of ∼1.6 km (Short, 2022). This spatial uncertainty has previously been addressed by aggregating datasets or extracting covariates from buffered areas around each ignition location (Pourmohamad et al., 2024). However, sensitivity analyses indicated that using spatially aggregated covariate data had little influence on model results (Fig. S2.1-S2.3). Reporting biases may also exist in fire occurrence data because fires occurring near roads and other infrastructure are more likely to be detected and documented (Aplet et al., 2026; Fusco et al., 2019). Nevertheless, we found no evidence that these reporting biases substantially influenced results at the scale of our study (Fig. S2.4). We obtained vegetation cover information from remote sensing estimates that contain inherent uncertainty (i.e., 6-14% root-mean-square error) (Allred et al., 2021), limiting our ability to identify precise relationships between vegetation cover and fire spread, including potential thresholds in flammability. Finally, we excluded the southern portion of the Sonoran Desert (primarily in the state of Sonora, Mexico) from our analyses because key spatial datasets (e.g., Allred et al., 2021; Short, 2022) were unavailable outside of the United States. The development of consistent, cross-border spatial datasets (e.g.,Villareal et al., 2026) is essential to expand analyses to the full geographic extent of the Sonoran Desert and improve our understanding of fire-regime change throughout this biodiverse region.

### 4.4. Implications for Fire-Regime Change in Drylands

A primary challenge in global change science is understanding and anticipating how interacting drivers can push ecosystems beyond their natural range of variation and into novel states (Turner & Seidl, 2023). Our results suggest that the Sonoran Desert is undergoing such a transition, as human-caused ignitions increasingly intersect with herbaceous fuels to support a novel fire regime. Anthropogenic activities are central to these changes in the Sonoran Desert and other dryland systems, where the human fingerprint on fire occurrence is often greater than in many other ecosystems (Balch et al., 2017; Fusco et al., 2016). However, different drivers influence different components of the fire regime. In the Sonoran Desert, human activities determine where fires start, herbaceous fuels determine whether they spread and become large, and climate variability influences both herbaceous fuel production and the potential for extreme fire behavior. Together, these processes create conditions that can facilitate rapid ecosystem transformation, threatening native species such as the iconic saguaro cactus (Aslan et al., 2021). Although climatic variables had relatively weak effects in our models, climate remains a critical driver of ecosystem transformation in drylands. It can influence the distribution of invasive plants (McMahon et al., 2021), the potential for recovery of native species (Shriver et al., 2018; Urza et al., 2021), and the ability of fires to ignite and spread (Abatzoglou et al., 2019). Thus, our findings in the Sonoran Desert provide a specific example of how interactions among people, fuels, and environmental conditions can drive fire-regime transformation across drylands worldwide, creating pathways to ecosystem change that become increasingly difficult to reverse once established (Brooks et al., 2004; D’Antonio & Vitousek, 1992; Fusco et al., 2022). Anticipating and managing these pathways is essential to sustain the biodiversity and ecological integrity of Earth’s dryland ecosystems.

## Acknowledgements

Our study area includes the traditional, ancestral lands of more than thirty Indigenous Nations (Native Land Digital, 2026). This research was supported by Joint Fire Science Program award #L25AC00100-1 and the Arizona Board of Regents, Technology and Research Innovation Fund (TRIF). Data processing and web application development were supported through an allocation of high-performance cloud computing resources provided by the Google Earth Engine Partner Tier program. Finally, we thank the participants of three practitioner workshops in Phoenix, Arizona, who provided valuable feedback on our analyses.

