## Supplementary Information for "Human-caused ignitions and plant invasion drive fire-regime change in a dryland biodiversity hotspot"

**Appendix S1**

*Description of Environmental Variables*

At each wildfire occurrence point (Qs 1 and 2) and within each fire perimeter (Q3), we acquired data describing the fractional cover of annual herbaceous cover (forbs and graminoids), bare ground, litter, perennial herbaceous cover, shrubs, and trees from the Rangeland Analysis Platform (RAP) v.3.0 dataset (Allred et al., 2021). These data provide annual, 30-m maps of each cover type, using Landsat imagery and convolutional neural networks, with estimated root-mean-square-error values ranging from 6.7-14.0%. We extracted these data at each fire occurrence point or the footprint of each fire perimeter (plus a 60-m outward buffer) for the year prior to fire occurrence.

We also obtained additional environmental variables that may influence fire behavior or suppression activity. First, we obtained a 10-m digital elevation model from the US Geological Survey (USGS) 3dep dataset (USGS, 2026). We used this DEM to calculate slope, terrain ruggedness, and topographic position. Terrain ruggedness represents the standard deviation of elevation in a surrounding area, and topographic position represents the difference between the elevation of a site and the average elevation of the surrounding area (Weiss, 2001). These metrics are sensitive to scale; we used a 910-m radius around each point, corresponding to the median daily spread rate of large fires in the western US (Coop et al., 2022). We extracted elevation, slope, terrain ruggedness, topographic position, and the continuous heat-insolation load index (Theobald et al., 2015) at the location of each fire occurrence point. We aggregated all layers to a 30-m resolution and aligned them to a common grid prior to extraction. Next, we extracted weather data around the timing of each fire ignition. Specifically, we obtained daily fire danger indices produced by the European Centre for Medium-Range Weather Forecasts (ECMWF) using fifth-generation (ERA5) reanalysis data at a 25-km spatial resolution and daily time step (Vitolo et al., 2020). We used Energy Release Component, Ignition Component, and Spread Component, three key indices used in the US National Fire Danger Rating System (NFDRS) to make fire management decisions. These are combined variables that incorporate daily weather, fuel moisture, and operational considerations, and, in comparison to individual metrics such as wind speed or air temperature, more directly tie to established fire suppression practices in the US.

As indicators of human use and infrastructure, we obtained the locations of major bodies of water, roads, and buildings within the study area. We obtained water data from a 90-m, global map of hydrography, restricting records to those representing major lakes or rivers (Yamazaki et al., 2019). We calculated the distance from the centroid of each 30-m pixel to the closest major lake or reservoir, as an indicator of potential recreational use. We acquired road data from the 2025 US Census Bureau TIGER line dataset (US Census Bureau, 2026). We separated road records into primary (MAF/TIGER feature class code [MTFCC] = S1100), secondary (MTFCC = S1200), and tertiary (MTFCC = S1400 or S1500) categories, representing different densities of traffic and vehicle use. For each category, we then calculated the Euclidean distance from the centroid of each 30-m pixel to the closest road line. We acquired the location of buildings from the Microsoft ML building footprint dataset (Microsoft, 2025). This dataset was developed from high-resolution imagery beginning in 2014, with continuous updates. We calculated the distance from the centroid of each 30-m pixel to the nearest building. Finally, we acquired transmission line data from Arderne et al. (2020). We calculated the distance from the centroid of each 30-m pixel to the nearest transmission line of any voltage.

**Appendix S2**

*Sensitivity of Random Forest Models to Spatial Resolution of Covariates*

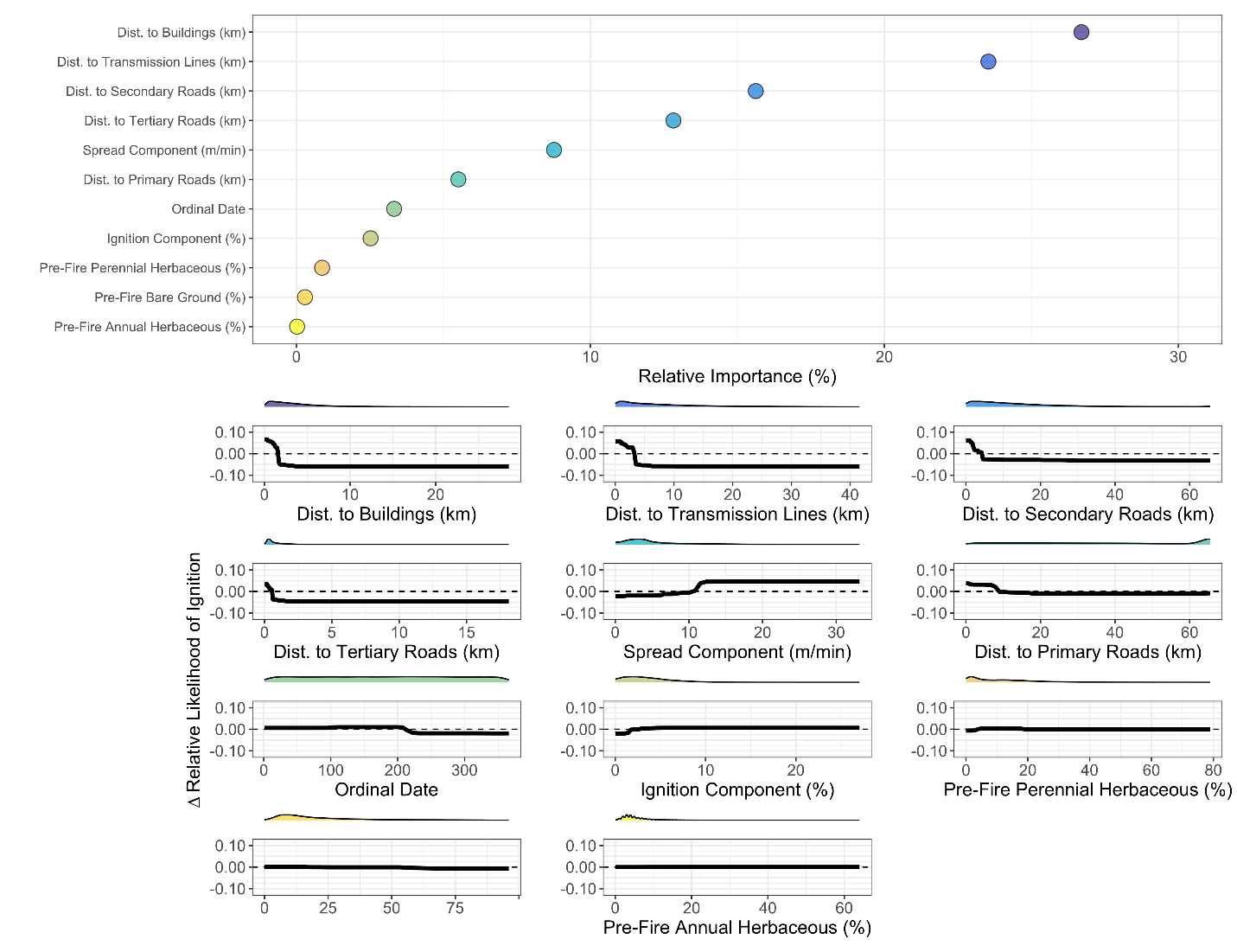

Figure S2.1: Results of the supplementary Random Forest model used to predict human ignition locations using spatial data at a 1.6-km spatial resolution, the maximum reported horizontal error of ignition locations in the FPA-FOD database (Short, 2022). Variables selected for inclusion, patterns of variable importance, and accumulated local effects plots were similar to those using higher-spatial-resolution covariates (Fig. 2 in main text). Likewise, out-of-bag accuracies were similar between this model (AUC = 0.857) and the model using higher-resolution covariates (AUC = 0.852).

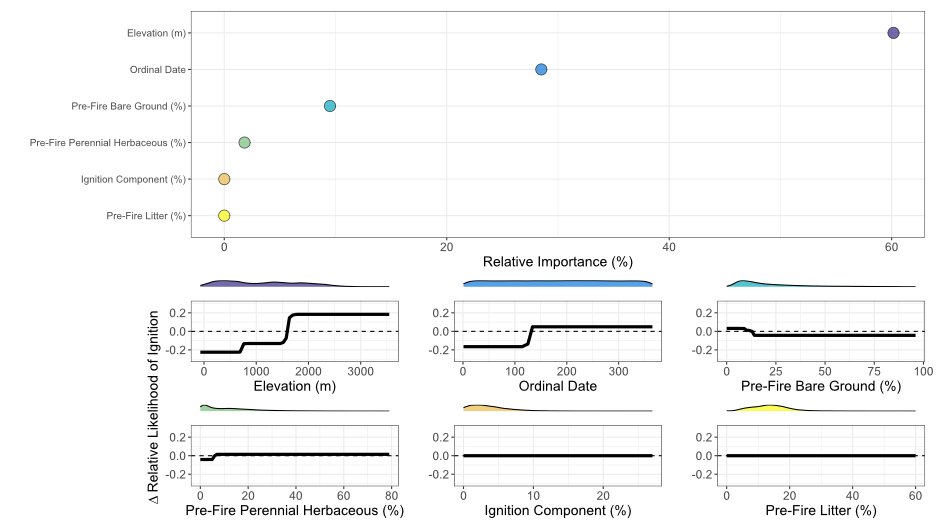

Figure S2.2: Results of the supplementary Random Forest model used to predict lightning-caused ignition locations using spatial data at a 1.6-km spatial resolution, the maximum reported horizontal error of ignition locations in the FPA-FOD database (Short, 2022). Variables included in model selection, variable importance, and patterns of accumulated local effects plots are similar to those using higher-spatial-resolution covariates (Fig. 3 in main text). Likewise, out-of-bag accuracies were similar between this model (AUC = 0.891) and the model using higher-resolution covariates (AUC = 0.909).

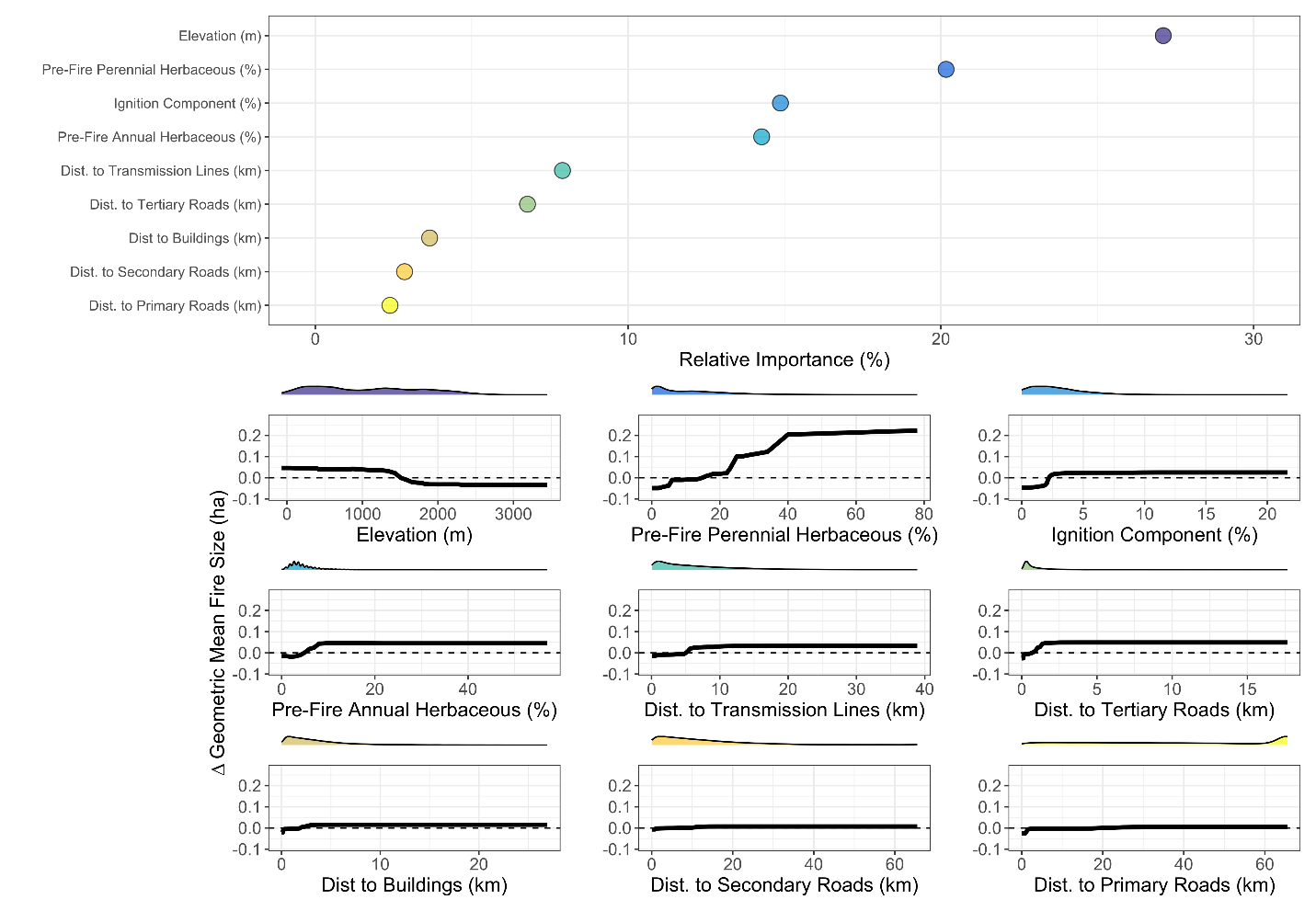

Figure S2.3: Results of the supplementary Random Forest model used to predict fire size using spatial data at a 1.6-km spatial resolution, the maximum reported horizontal error of ignition locations in the FPA-FOD database (Short, 2022). Variables included in model selection, variable importance, and patterns of accumulated local effects plots are similar to those using higher-spatial-resolution covariates (Fig. 4 in main text). Likewise, out-of-bag accuracies were similar between this model (Pearson’s R = 0.338) and the model using higher-resolution covariates (Pearson’s R = 0.339).

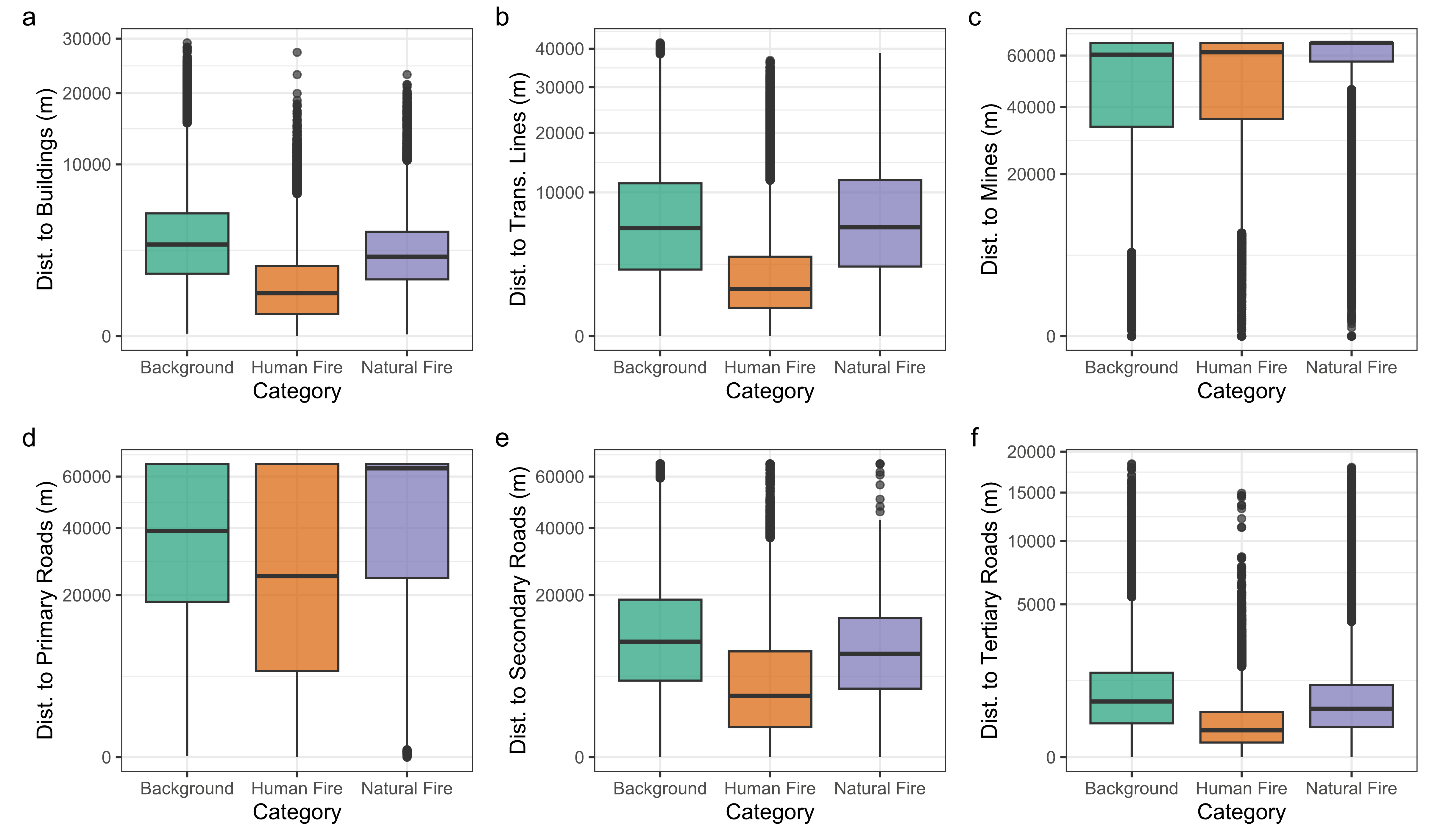

Figure S2.4: An assessment of potential reporting biases in agency records (Short, 2022), where we assumed that human or lightning-caused ignitions may be more likely to be reported near roads or human infrastructure (Aplet et al., 2026; Fusco et al., 2019). However, we found that distances between lightning-caused ignitions and human infrastructure did not meaningfully differ from the background distribution across the study area, whereas human-caused fires were typically closer to infrastructure. Because potential reporting biases would be likely to be present across both human and lightning ignitions (e.g., both categories would be more likely to be near homes or roads than background points), this suggests that reporting biases did not play a strong role in our human ignition models.
